# TRMT2B is responsible for both tRNA and rRNA m^5^U-methylation in human mitochondria

**DOI:** 10.1101/797472

**Authors:** Christopher A. Powell, Michal Minczuk

## Abstract

RNA species play host to a plethora of post-transcriptional modifications which together make up the epitranscriptome. 5-methyluridine (m^5^U) is one of the most common modifications made to cellular RNA, where it is found almost ubiquitously in bacterial and eukaryotic cytosolic tRNAs at position 54. Here, we demonstrate that m^5^U54 in human mitochondrial tRNAs is catalysed by the nuclear-encoded enzyme TRMT2B, and that its repertoire of substrates is expanded to ribosomal RNAs, catalysing m^5^U429 in 12S rRNA. We show that TRMT2B is not essential for viability in human cells and that knocking-out the gene shows no obvious phenotype with regards to RNA stability, mitochondrial translation, or cellular growth.

## Introduction

The accurate expression of the human mitochondrial genome (mtDNA) is essential for the faithful synthesis of 13 components of the oxidative phosphorylation machinery. In order to achieve this, the 22 tRNAs and 2 rRNAs encoded in the mtDNA undergo significant post-transcriptional modification performed by nuclear-encoded proteins imported into mitochondria^1^. Recent data indicate that defects in the nucleotide modification of mitochondrial (mt-) RNA can frequently lead to human disorders of mitochondrial respiration^2–5^

5-methyluridine (m^5^U), is found with high occurrence in bacterial and eukaryotic tRNAs at position 54 in the T-loop, including, although significantly less common, human mitochondrial tRNAs. Although first identified in tRNAs, the T-loop motif has now been identified in a wide array of different non-coding RNAs, including the tmRNA of bacteria^6^, RNase P RNA^7^, self-splicing introns^8^, and riboswitches^9^, implying that this frequently occurring motif is an important structural building block. T-loops have additionally been implicated in stabilising the tertiary fold of rRNA, this has been suggested to occur either via base pairing/base pairing tertiary interactions or by stabilizing stems by forming a non-interacting caps^10^. The m^5^U modification is also found in a number of rRNA species, including that of the mitochondrial small rRNA in hamster^11,12^ however as before, their exact contribution to the surrounding tertiary structure is uncertain.

The vast majority of methyltransferases that catalyse m^5^U formation utilise SAM (S-adenosyl-methionine) as a methyl donor, however a small number including TrmFO from *B*. *subtilis*^13^ and RlmFO from *M*. *capricolum*^14^, use 5, 10-methylenetetrahydrofolate (M-THF) as a methyl source. The most well studied m^5^U-methyltransferases, all of which fall into the former, SAM-dependent, group, are the three expressed by *E*. *coli*: RlmC, RlmD, and TrmA. Both RlmC and RlmD methylate the 23S rRNA of the large subunit, with the former producing m^5^U747 and the latter producing m^5^U1939^15^. The aforementioned m^5^U54 tRNA modification is catalysed by TrmA, which also methylates the T-loop in *E.coli* tmRNA^16^. TrmA has additionally been found to methylate bacterial 16S rRNA *in vitro*, however this modification is not found *in vivo*^17^. The proposed catalytic mechanism utilised by m^5^U-methyltransferases has been supported by the crystal structure of the TrmA-tRNA complex in which the sulphhydryl-group of a conserved cysteine acts as a nucleophile to attack C6, leading to the formation of a covalent bond between the two. The resulting negative charge on O4 prompts a second nucleophilic attack by the C5 carbon on the methyl group of SAM, forming m^5^U. A second highly conserved residue, a glutamate, acts as a base to abstract a proton from C5 leading to the elimination of the covalent adduct and its release from the regenerated active site^18^. The identification and characterisation of TrmA has not brought the field significantly closer to understanding the exact role of m^5^U54, as mutations in *E.coli* TrmA that lead to the complete loss of methyltransferase activity revealed no perceptible difference in growth rate, codon recognition, ribosome interaction, or translation rate *in vivo*^19^. A growth defect in the TrmA mutant cells was only observed following growth in a mixed culture in which the two strains directly competed. Interestingly, the TrmA protein itself, but not its known enzymatic activity, is found to be essential for viability. An insertion within the *trmA* gene corresponding to the N-terminus of the protein is lethal, therefore it is proposed that TrmA has a secondary essential function that is separate from its methyltransferase activity^20^.

The loss of m^5^U54 from both cytoplasmic and mitochondrial tRNAs was first observed in eukaryotic cells following the deletion of the *trm2* gene in *S.cerevisiae*^21^. As was the case for bacterial TrmA, the loss of Trm2 showed no physiological defect compared to wild-type cells, however even in cocultures of mixed populations, the wild-type cells demonstrated no selective advantage over the mutant strain after 35 generations. The requirement for trm2 is further contrasted from that of trmA by the nonessential nature of not only m^5^U54 but of the protein itself, as a complete deletion of *trm2* also shows no physiological defect^22^. The large majority of homology between TrmA and Trm2 lies towards the C-terminus containing the known methyltransferase domain, with little homology towards the N-terminus. This, therefore, supports the notion that the unidentified essential function of TrmA is situated towards the N-terminus, and that this function is either not performed by Trm2 or there is a greater degree of functional redundancy in eukaryotic cells. Although the absence of Trm2 in isolation shows no physiological effect, its deletion has been demonstrated to induce lethality in four strains carrying mutations in tRNA^Ser(CGA)23^, indicating that the stabilising nature of m^5^U54 may only be observed when the structure of a tRNA is affected in another manner. Intriguingly, although three of the four mutants require catalytically active Trm2, and therefore m^5^U54, to be present for stability, one mutation described is entirely rescued through the expression of a catalytically inactive Trm2. It is therefore suggested that in addition to introducing the stabilising role of m^5^U54, Trm2 itself may possess a chaperone-like function that is separate from its catalytic role. A separate endo-exonuclease activity has been claimed for Trm2, which is preserved following deletion of the C-terminal methyltransferase domain^24^. Through this activity, Trm2 is proposed to contribute to the repair of double-strand breaks by the 5’-3’ resection of DNA ends. However, how this additional activity would explain the phenotypes observed with tRNA^Ser(CGA)^ mutants is unclear.

Efforts to characterise the modification profile of human tRNAs have identified the presence of m^5^U54 in the majority of cytoplasmic tRNAs as well as a small number of mitochondrial tRNAs. A very recent study demonstrated that human TRMT2A is responsible for introduction of the majority of m^5^U in human RNA, and that it targets U54 of cytosolic tRNAs^25^ However, the enzyme responsible for m^5^U in human mt-tRNA remains to be characterised. Likewise, the methyltransferase responsible for the introduction of m^5^U429 in mammalian mitochondrial 12S rRNA is yet to be identified. In this work, we aim to identify the ortholog of Trm2 operating in human mitochondria and to study the consequences of its loss.

## Results

### TRMT2B is a putative methyltransferase localised to human mitochondria

The m^5^U54 modification has been identified in both cytoplasmic and mitochondrial tRNAs in humans^26^, with the former recently identified to be catalysed by TRMT2A^25^, one of the two human orthologs of yeast Trm2 along with TRMT2B. As a mitochondrial role for TRMT2A was not ruled out by Carter *et al*.^25^, we set out to identify which of these paralogs may be operating within human mitochondria. An alignment of protein sequences from a range of species shows that the SAM binding site is highly conserved in the TRMT2A and TRMT2B paralogs. Whilst this is also the case for the remainder of the active site in TRMT2A, TRMT2B displays some dramatic divergences in some species (**Figure 1**). The proposed common mechanism for SAM-dependent m^5^U methyltransferases, supported by the crystal structure of TrmA^18^, entirely depends on the involvement of a nucleophilic cysteine and a proton extracting glutamate (**Figure S1**). Whilst these two residues are conserved in all species analysed for TRMT2A, a number of species contain mutations altering the catalytic cysteine in TRMT2B, which would be predicted to very severely impede the enzymes methyltransferase activity. The majority of these species belong to a single subfamily, the Bovinae, in which this otherwise well conserved cysteine is substituted for tyrosine (**Figure 1**). The finding that the bovine TRMT2B homolog appears catalytically inactive is in agreement with the initially surprising observation that m^5^U54, one of the most common tRNA modifications, is absent from all bovine mt-tRNAs^27^. However, m^5^U54 is highly abundant in bovine cyto-tRNAs^26^, it is tempting to speculate therefore, that these cytosolic substrates are methylated by the functional TRMT2A, which does not operate on mt-tRNAs.

**Figure 1.**
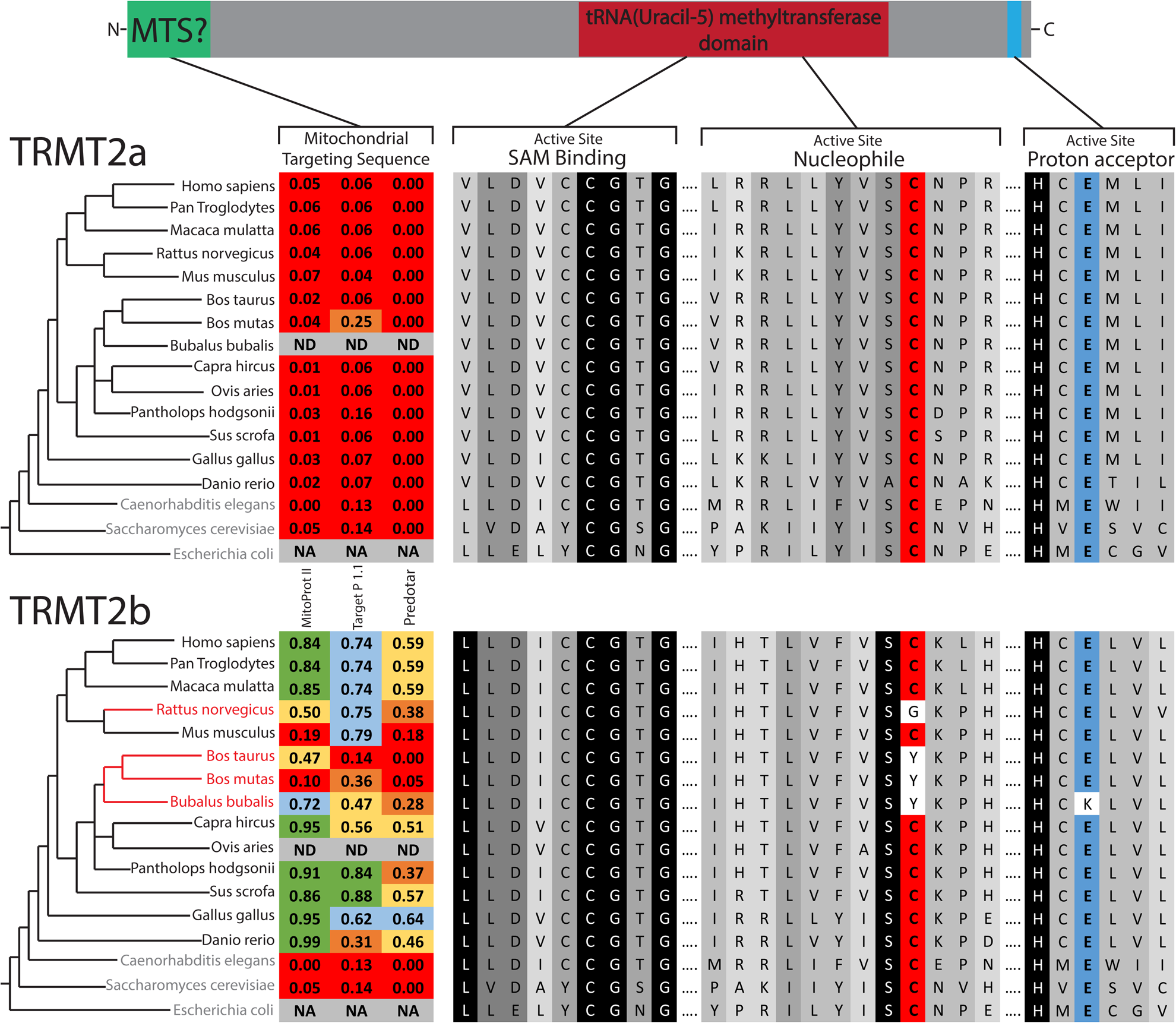
Sequence Analysis of m^5^U54-methyltransferase homologs. Schematic of protein domains in both TRMT2A and TRMT2B. Homologous sequences identified through BLAST searches aligned for key catalytic regions, with the degree of shadowing representing the extent of conservation for a given residue. Nucleophilic cysteine shown in red, Proton extracting glutamate shown in blue. Species that contain only a single predicted m^5^U54-methyltransferase indicated in blue text. Species that contain a TRMT2B sequence that is predicted to be catalytically inactive are indicated in red text. These two paralogs were analysed for the presence of an N-terminal mitochondrial localisation signal (MTS) using computational prediction tools. Higher scores (between 0 and 1) indicate a higher probability of mitochondrial localisation, which are also colour coded. NA, not applicable; ND, no data

The observation that bovine cyto-tRNAs are methylated at U54, whilst mt-tRNAs are not, could be explained by the solely cytosolic localisation of a catalytically active TRMT2A, and the mitochondrial localisation of a catalytically inactive TRMT2B. To test this hypothesis, the sequences of both paralogs were analysed for the likelihood of an N-terminal mitochondrial targeting sequence (MTS). In all three of the programs used to predict mitochondrial localisation (MitoProt II^28^, Target P 1.1^29^, and Predotar^30^), the TRMT2A sequence returned very low scores across all species analysed, consistent with a non-mitochondrial role. These scores were significantly higher for TRMT2B, indicating possible mitochondrial targeting (**Figure 1**). To confirm the mitochondrial localization of human TRMT2B, HeLa cells were transiently transfected with a C-terminal GFP-tagged TRMT2B and localised based on fluorescence. Through this analysis, TRMT2b was found to colocalise with the known mitochondrial protein TOM20 (**Figure 2**). Taken together these results show TRMT2B is a mitochondria-localised protein.

**Figure 2.**
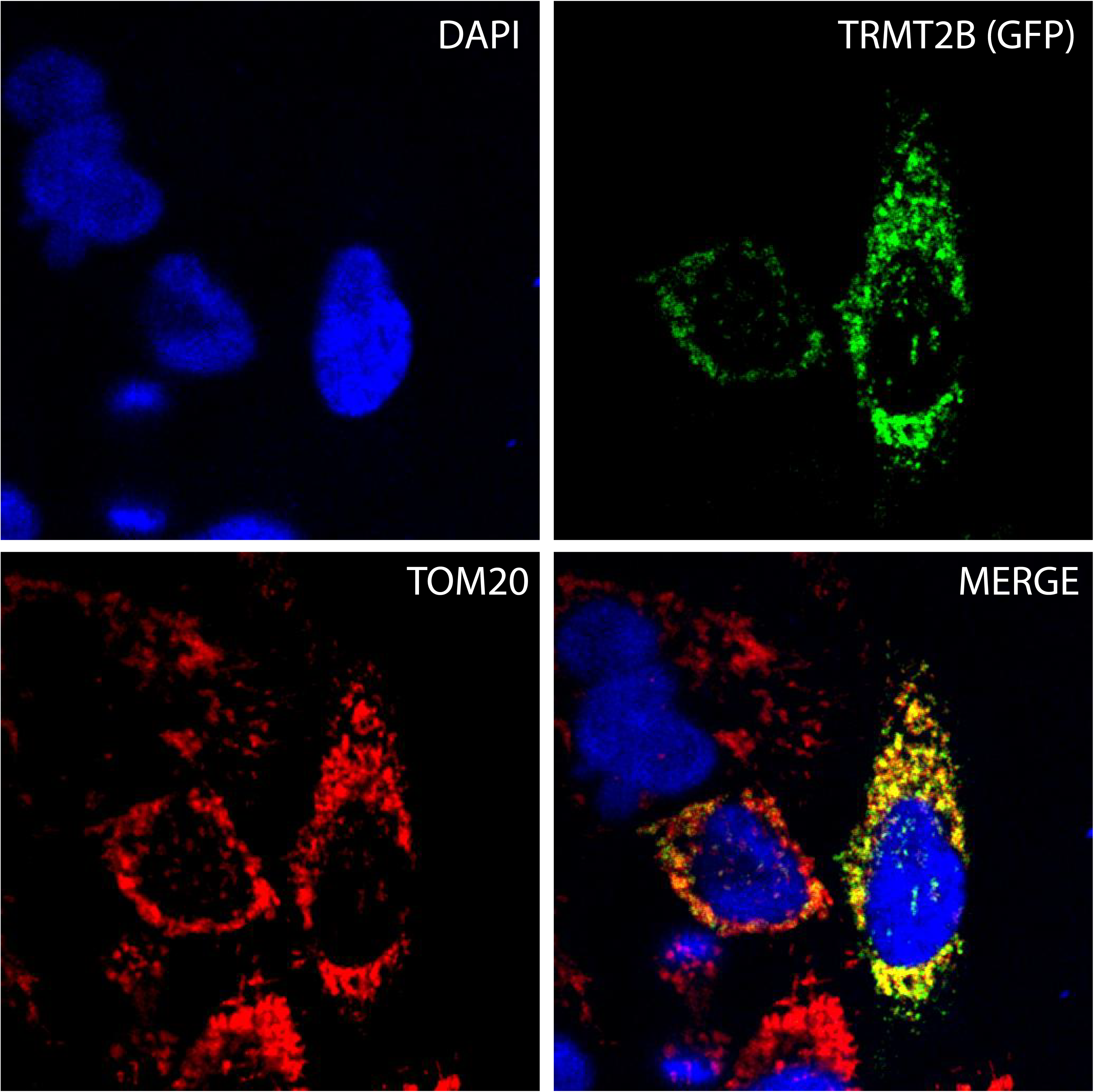
TRMT2B is localised to mitochondria. TRMT2B-GFP cDNA construct transfected into HeLa cells and detected by fluorescence (top right, green). Nuclei were stained using DAPI (top left, blue). The mitochondrial network was stained using antibodies against the known mitochondrial protein TOM20 (bottom left, red). Colocalisation of TRMT2B and TOM20 appears as yellow in a digitally overlaid image (bottom right).

### TRMT2B is a mitochondrial m^5^ U54 tRNA methyltransferase

Next, we set out to determine mtRNA targets for TRMT2B. Due to the location of position 5 on the Hoogsteen, rather than Watson-Crick edge (**Figure S2A**), its methylation to form m^5^U does not interfere with the processivity of a reverse transcriptase, and therefore cannot be detected by primer extension without prior treatment. The presence of m^5^U can be inferred, however, due to its resistance to depyrimidination in the presence of hydrazine, a process which unmodified uracil is susceptible to (**Figure S2A**). Subsequent treatment with aniline results in phosphodiester bond cleavage at abasic sites, which are then detected due to the accumulation of primers that are unable to extend further (**Figure S2B**). In order to test the predicted role of TRMT2B, the aforementioned technique was used to test the U54 methylation status following transfection with siRNAs targeted to deplete TRMT2B. For mt-tRNA^Pro^, which has been previously identified as containing m^5^U54^31^, the degree of stalling at U54, relative to downstream U49, is notably increased following TRMT2B siRNA transfection compared to an untransfected control (**Figure 3A**). To confirm the role of TRMT2B in m^5^U54 methylation the same assay was repeated using RNA extracted from a TRMT2B knock-out HAP1 cell line. Correspondingly, the degree of stalling at U54 was found to be drastically increased in the knock out cell line relative to a control, consistent with the absence of m^5^U54 (**Figure 3B**). In agreement with the previously identified role for TRMT2A^25^, the m^5^U54 status of cytosolic tRNAs were found to be unaffected following the loss of TRMT2B activity (**Figure S3**). Collectively, these results show TRMT2B is a mitochondrial protein responsible for the modification of position 54 in mt-tRNAs.

**Figure 3.**
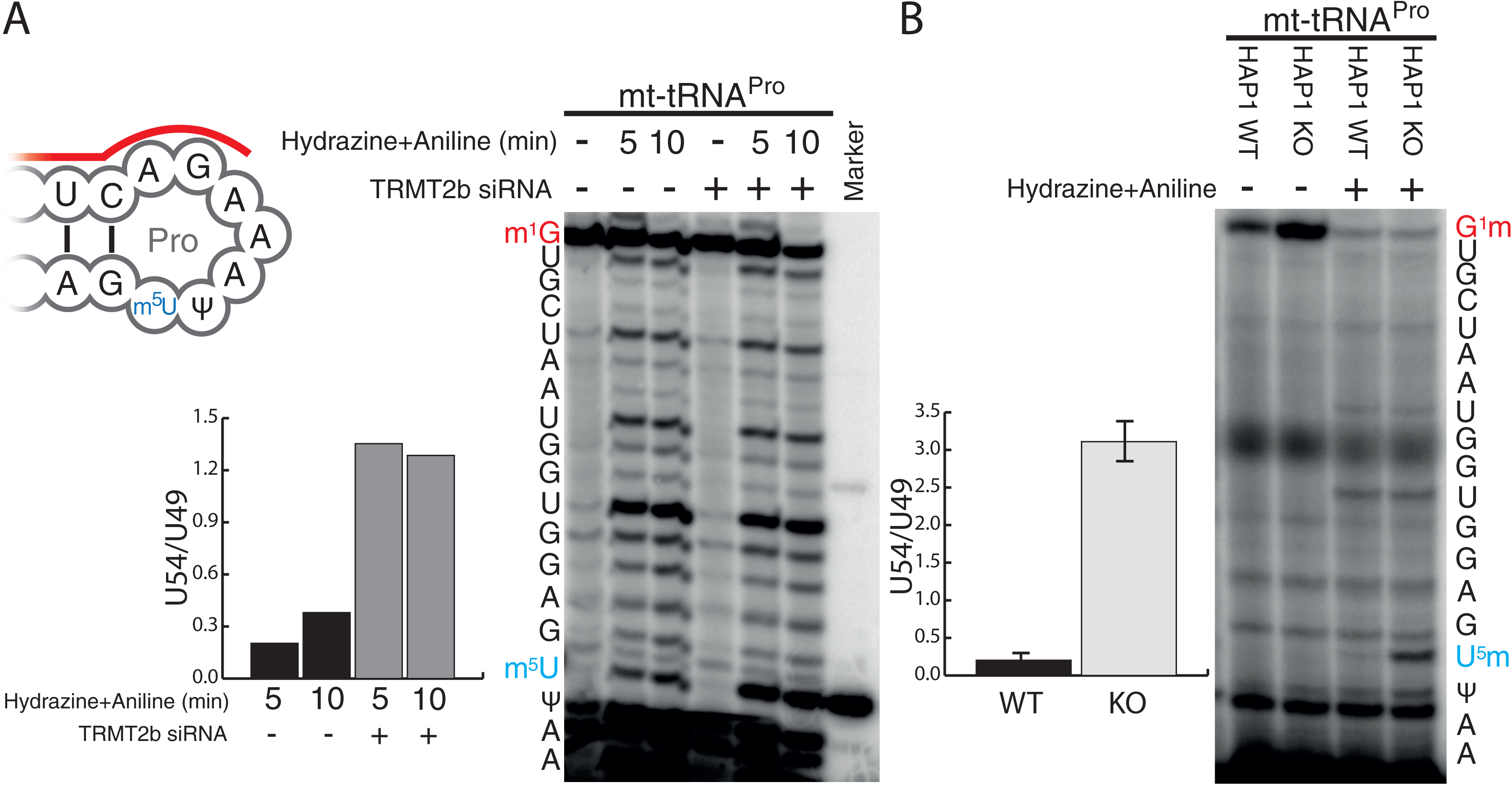
TRMT2B catalyses m^5^U54 in mt-tRNA^Pro^. **(a)** Schematic of mt-tRNAPro T-loop showing annealed primer to be extended (red line) and the position of m^5^U54 (blue text). HeLa cell derived RNA, either following a 6-day siRNA mediated depletion of TRMT2B or untreated, was subsequently either untreated (-) or treated with hydrazine for either 5 or 10 minutes, followed by aniline, to specifically cleave at unmodified uridine residues. This RNA was subjected to RT-PEx using a [^32^P]-end labelled primer complementary to the region upstream of m^5^U54 (red line). The nucleotide sequence of the tRNA, corresponding to stalling events at each position, is shown to the side of the panel. Quantification values represent the ratio between stalling at U54 and stalling at the next uridine residue (U49), after the values in the untreated lanes had been subtracted from both to account for background. **(b)** RT-PEx reactions as performed above with RNA derived from a HAP1 parental cell line with wild-type (WT) or a HAP1 TRMT2B knockout cell line (KO). Error bars = SEM, n = 3.

### TRMT2B is a mitochondrial m^5^ U429 12S rRNA methyltransferase

An m^5^U methylation has also been previously identified at U426 within the small 12S rRNA from hamster mitochondria^11^. A sequence alignment of this region shows a very high degree of conservation across a range of eukaryotic species (**Figure 4A**). This sequence conservation is also present in the small 16S rRNA from *E.coli* where it forms the 790-loop, and whilst the corresponding uracil (U788) is not methylated, mutations within this loop yielded no fully formed 30S subunit^32^. The corresponding human residue, U429, is located within the central domain of 12S rRNA in helix 27 (**Figure 4B**), the loop of which is in very close proximity to the codon-anticodon site between bound mRNA and tRNA (**Figure 4C**). As this loop displays similarity to the consensus T-loop, and m^5^U has been detected at this position in hamster mitochondria, 12S U429 may also act as a substrate for TRMT2B, in addition to mt-tRNAs. To test whether TRMT2B is responsible for modification of 12S U429 in human mitochondria, the presence or absence of m^5^U was tested following an RNA treatment of hydrazine and aniline in wild type and TRMT2B knockout cells. Very little stalling at U429 was observed in wild type cells (**Figure 4D**), consistent with a modification conferring resistance to hydrazine. On the contrary, stalling at U429 is significantly increased in RNA extracted from TRMT2B knockout cells(**Figure 4D**). These results are consistent with human 12S containing an m^5^U methylation, as has been shown for hamster, and with this methylation being catalysed by TRMT2B.

**Figure 4.**
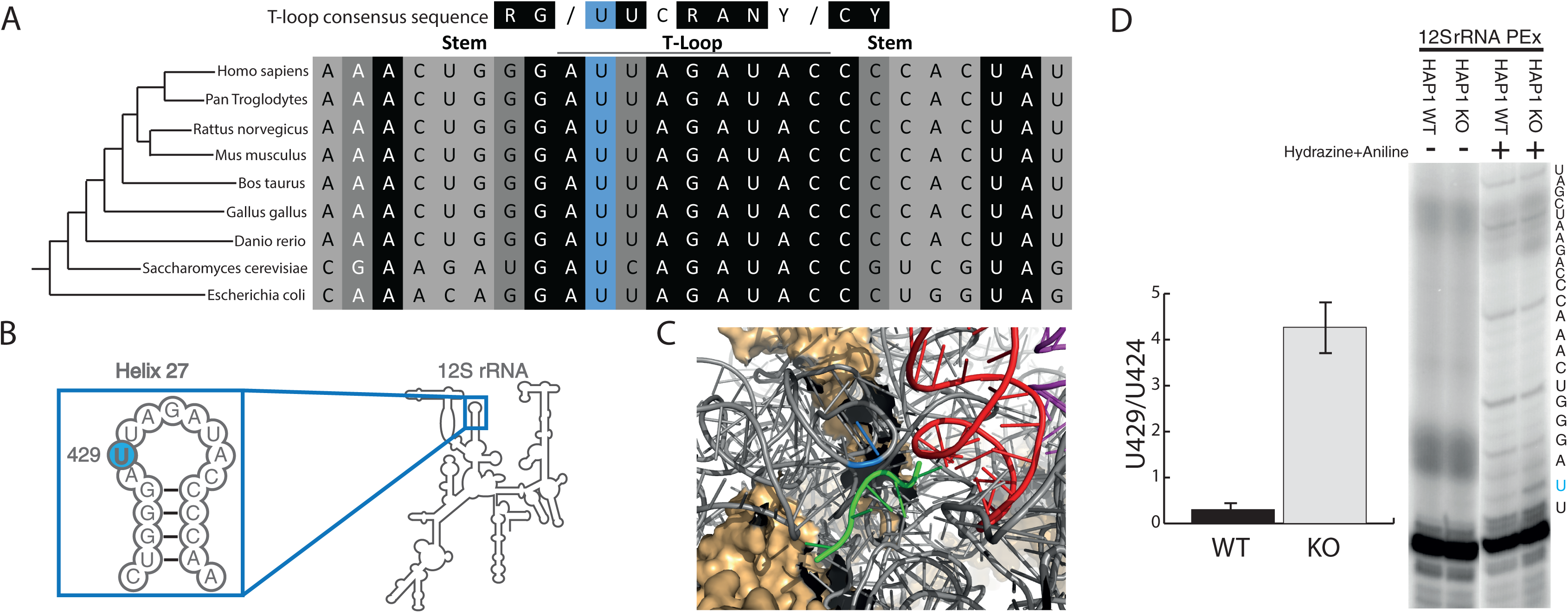
TRMT2B catalyses m^5^U429 in 12S mitochondrial rRNA. **(a)** Alignment of the small ribosomal RNA (rRNA) from a range of species in a region corresponding to helix 27 in human 12Smt-rRNA. The degree of shadowing represents the extent of conservation for a given residue, with U429 (or the corresponding position in other species) shown in blue background. The T-loop consensus sequence is displayed above for comparison. **(b)** Schematic of human 12S mt-rRNA, helix 27, and the location of m^5^U429 (blue circle). **(c)** The structure of the human mitoribosome highlighting U429 (blue), the bound mRNA (green), and the adjacently bound tRNA (red). **(d)** Separation and detection of RT-PEx products using a [^32^P]-end labelled primer complementary to the region upstream of m^5^U429 in 12S rRNA. Extension reactions performed on RNA derived from a HAP1 wild-type (WT) and a HAP1 TRMT2B knockout cell line (KO), with or without hydrazine-aniline treatment. The nucleotide sequence of 12S rRNA is shown to the side of the panel. Quantification values represent the ratio between stalling at U429 and stalling at the next uridine residue (U424). Error bars = SEM, n = 4.

### Mitochondrial tRNA stability and aminoacylation are unaffected by the loss of TRMT2B

Incorrect folding as a consequence of hypomodification has been shown to disrupt the interaction between a tRNA and its cognate aminoacyl-tRNA synthetase^33,34^. In order to assess the impact of m^5^U54 loss on tRNA charging, RNA extracted from WT and TRMT2B KO cells was subjected to acidic-PAGE, allowing for charged and uncharged tRNAs to be distinguished by their differing electrophoretic mobilities. In this analysis, the degree of aminoacylation for three m^5^U54-containing mt-tRNAsshows no change as a result of the loss of TRMT2B (**Figure 5A**). Interestingly, the migration pattern of mt-tRNA^LeuUUR^ is altered between the wild-type and knockout cell lines, but this shift is present in both the acylated and deacylated samples, and therefore independent of aminoacylation. The conditions in which acidic-PAGE gels are run may allow for the preservation of certain tRNA secondary structures^35^, and therefore this electrophoretic shift is likely an indication of the structural contribution made by m^5^U54.

**Figure 5.**
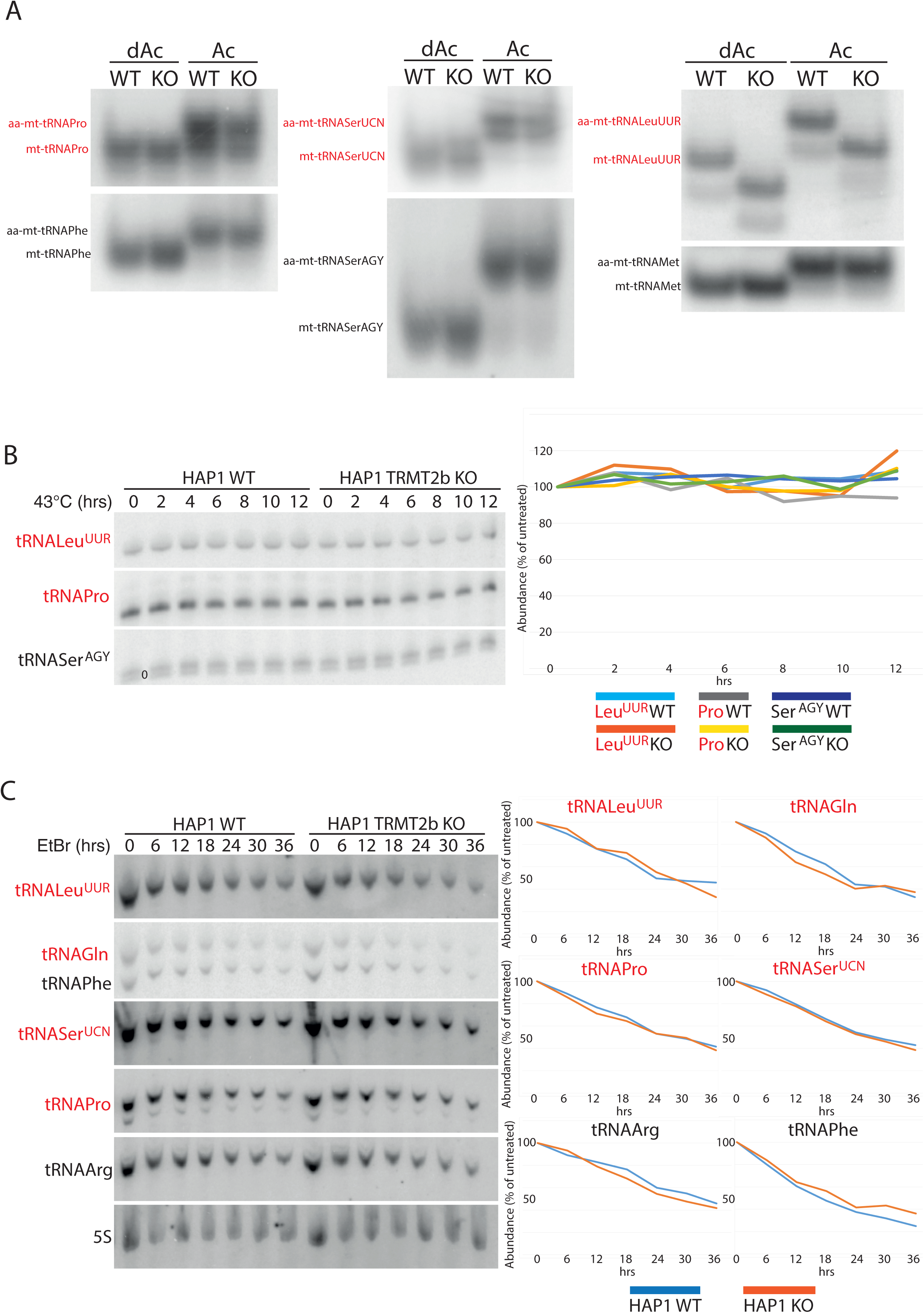
tRNA aminoacylation and stability following the loss of TRMT2B. **(a)** High resolution acidic polyacylamide gel electrophoresis (PAGE) northern blot analysis of total RNA extracted from HAP1 wild-type (WT) or TRMT2B knockout cell line (KO). RNA was either maintained in low pH conditions to preserve aminoacylation (Ac), or intentionally deacylated prior to being run on the gel (dAc). The blots were probed with the mt-tRNA-specific riboprobes as indicated, with the tRNAs known to contain m^5^U54 in red text. **(b)** High resolution PAGE northern blot analysis of total RNA extracted from HAP1 parental cell line (WT), or a HAP1 TRMT2B knockout cell line (KO), following their growth at 43 °C for between 2 and 12 hours. Control cells kept at 37 °C shown as ‘0 hrs’. The blots were probed with the mt-tRNA-specific riboprobes as indicated, with the tRNAs known to contain m^5^U54 in red text. Quantification values were plotted as a percentage of the untreated 0 hr RNA. **(c)** As above following exposure to 250 μg/mL ethidium bromide (EtBr) for between 6 and 36 hours. Control cells grown in media containing no EtBr shown as ‘0 hrs’. The blots were probed with the mt-tRNA-specific riboprobes as indicated, with the tRNAs known to contain m^5^U54 in red text. Quantification values were plotted as a percentage of the untreated 0 hr RNA, after they had been normalised to 5S to control for RNA loading.

As *in vitro* studies have pointed towards a role for m^5^U54 in tRNA thermal stability^36^, the loss of TRMT2B has the potential to induce a temperature-sensitive phenotype. Whilst the stability of mature tRNAs is typically unaffected by heat-stress, a 6 hour heat-shock at 43 °C induces a 60 % reduction in the levels of initiator cyto-tRNA^Met^, which coincidentally is one of the few cytoplasmic tRNAs to lack m^5^U54^37^. The steady state levels of mt-tRNAs were assessed through northern blotting following RNA extraction from wild-type and knockout cells which were incubated at 43 °C for between 2 and 12 hours, or kept at 37 °C (0 hrs). No decrease in the steady state levels was observed for m^5^U54-bearing or m^5^U54-absent tRNAs, either in the wild-type or the knock-out cells (**Figure 5B**). Whilst this suggests no decrease in the stability of m^5^U54-lacking tRNAs, a constant steady state level could be maintained by a mtDNA transcription rate sufficiently high to replenish any degraded tRNA. Therefore, in order to test if this is the case, cells were treated with ethidium bromide to block mitochondrial transcription^38,39^. RNA was extracted from wild-type and TRMT2B knockout cells at a range of time points between 6 and 36 hours, following the addition of 250 μg/mL of ethidium bromide, along with untreated cells (0 hrs). The rate of degradation for m^5^U54-bearing or m^5^U54-absent tRNAs showed no variation between the wild-type and knockout cell line (**Figure 5C**). Therefore, these data suggest no difference in the stability of mt-tRNAs following the loss of m^5^U54.

### Mitoribosomestability is not disrupted by the loss of TRMT2B

The disruption of assembly pathways through the loss of an rRNA modification has been previously shown to result in the degradation of subunits that are unable to be incorporated into monosomes^40^. In order to determine if TRMT2B activity contributes towards mitoribosome biogenesis, the steady state levels of protein components of the mitochondrial small 28S subunit were, therefore, assessed by western blotting. However, of the three MRPS proteins determined, the loss of m^5^U429 appears to have no discernible effect on their steady state levels (**Figure 6A**). Likewise, the loss of m^5^U429 also has no effect on the steady-state levels of 12S mt-rRNA itself relative to 16S mt-rRNA (**Figure 6B**). As for mt-tRNAs, the effect of m^5^U429 on 12S thermal stability was determined by a 43 °C heat shock for between 2 and 12 hours. Interestingly, 12S mt-rRNA displayed a significantly greater susceptibility to heat-induced degradation compared to 16S rRNA, however, no difference in the rate of degradation was observed between the wild-type and knockout cell lines (**Figure 6C**). An analysis of 12S stability in the absence of replenishment from mitochondrial transcription also identifies no difference in the stability of 12S mt-rRNA resulting from the loss of TRMT2B (**Figure 6D**).

**Figure 6.**
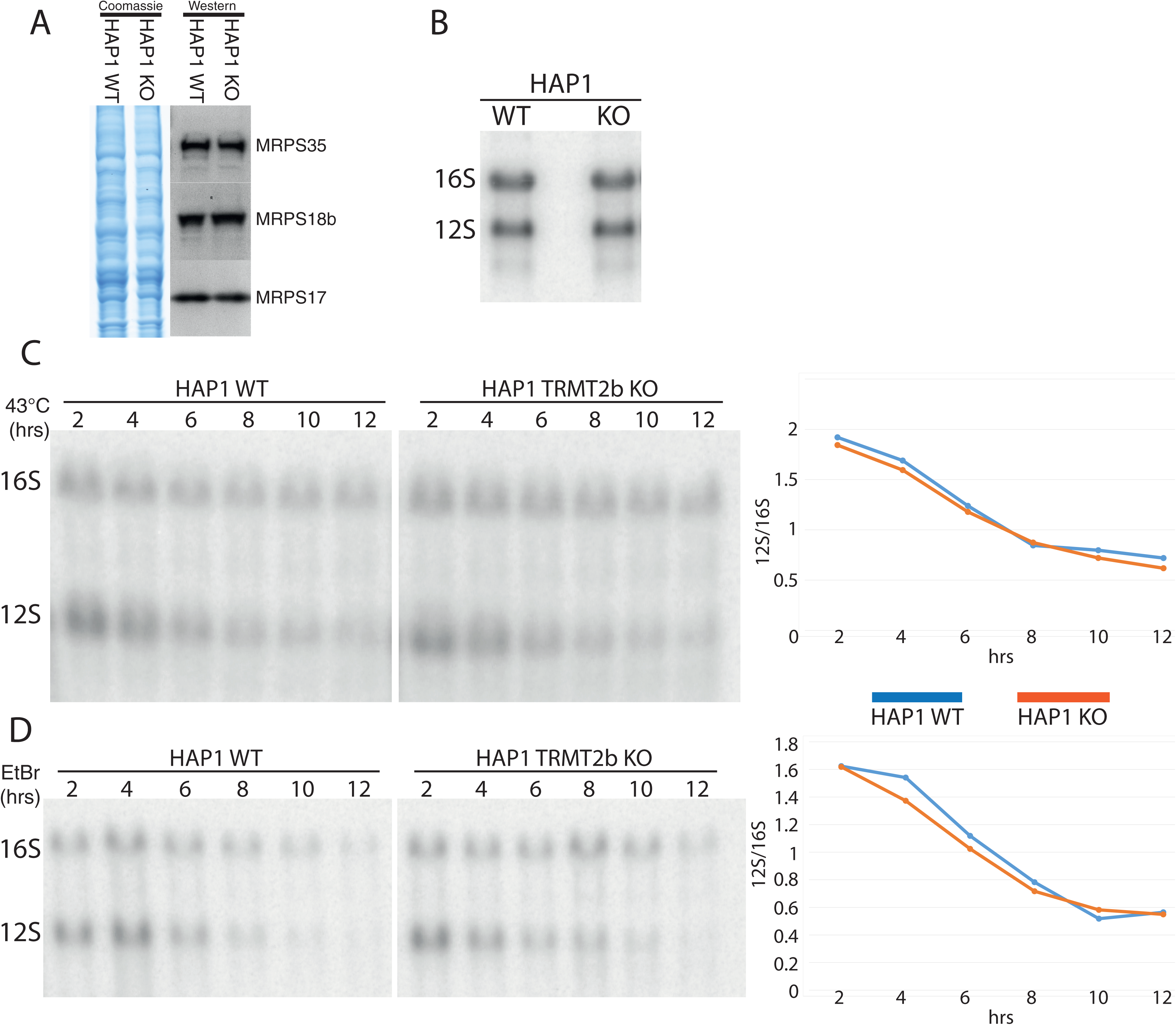
Mitoribosome integrity following the loss of TRMT2B. **(a)** Western blot using lysates derived from HAP1 WT and KO probed using antibodies against three protein components on the small subunit. Coomassie gel staining shown as loading control. **(b)** Northern blot analysis of total RNA extracted from HAP1 parental cell line (WT), or a HAP1 TRMT2B knockout cell line (KO). **(c)** Northern blot analysis of total RNA extracted from HAP1 WT and TRMT2B KO cell lines following their growth at 43 °C for between 2 and 12 hours. Quantification values plotted as the ratio between 12S and 16S. **(d)** Northern blot analysis of total RNA extracted from HAP1 WT and TRMT2B KO cell lines following their exposure to 250 μg/mL ethidium bromide (EtBr) for between 2 and 12 hours.

### The loss of TRMT2B has no significant impact on mitochondrial translation

Although the loss either m^5^U54 or m^5^U429 have caused no detectable differences in the stability of mt-tRNAs or 12S mt-rRNA, their absence may instead be detrimental to the efficiency or fidelity of mitochondrial translation. To this end, the steady state levels of representative subunits from respiratory chain complexes were assessed through western blotting, however this displayed no significant difference resulting from the loss of TRMT2B (**Figure 7A**). Similarly, the rate of mitochondrial protein synthesis was measured by the incorporation of [^35^S] labelled methionine into nascent polypeptides. Once again, no difference between the wildtype and knockout cell line was observed (**Figure 7B**). Potentially, differences in the mitochondrial translation rate that are too subtle to distinguish through the assays described above, would be expected to provide cells with a growth advantage that would become apparent over a significantly long period of expansion. Wildtype and TRMT2B knockout cells were plated at a low confluency and the cell density measured over a 140 hour time course of growth. These growth curves were performed in either a high glucose medium or low glucose medium, with the latter exacerbating any growth disadvantage stemming from mitochondrial dysfunction, as cells are forced to rely more greatly on oxidative phosphorylation than glycolysis. Of note, growth in medium containing galactose as the sole carbon source, resulted in considerable cell death in both the wildtype and TRMT2B knockout HAP1 cell lines, obscuring accurate measurements. Under both conditions tested, the TRMT2B knockout cell line exhibited no statistically significant decrease in growth rate compared with the wildtype control (**Figure 7C**).

**Figure 7.**
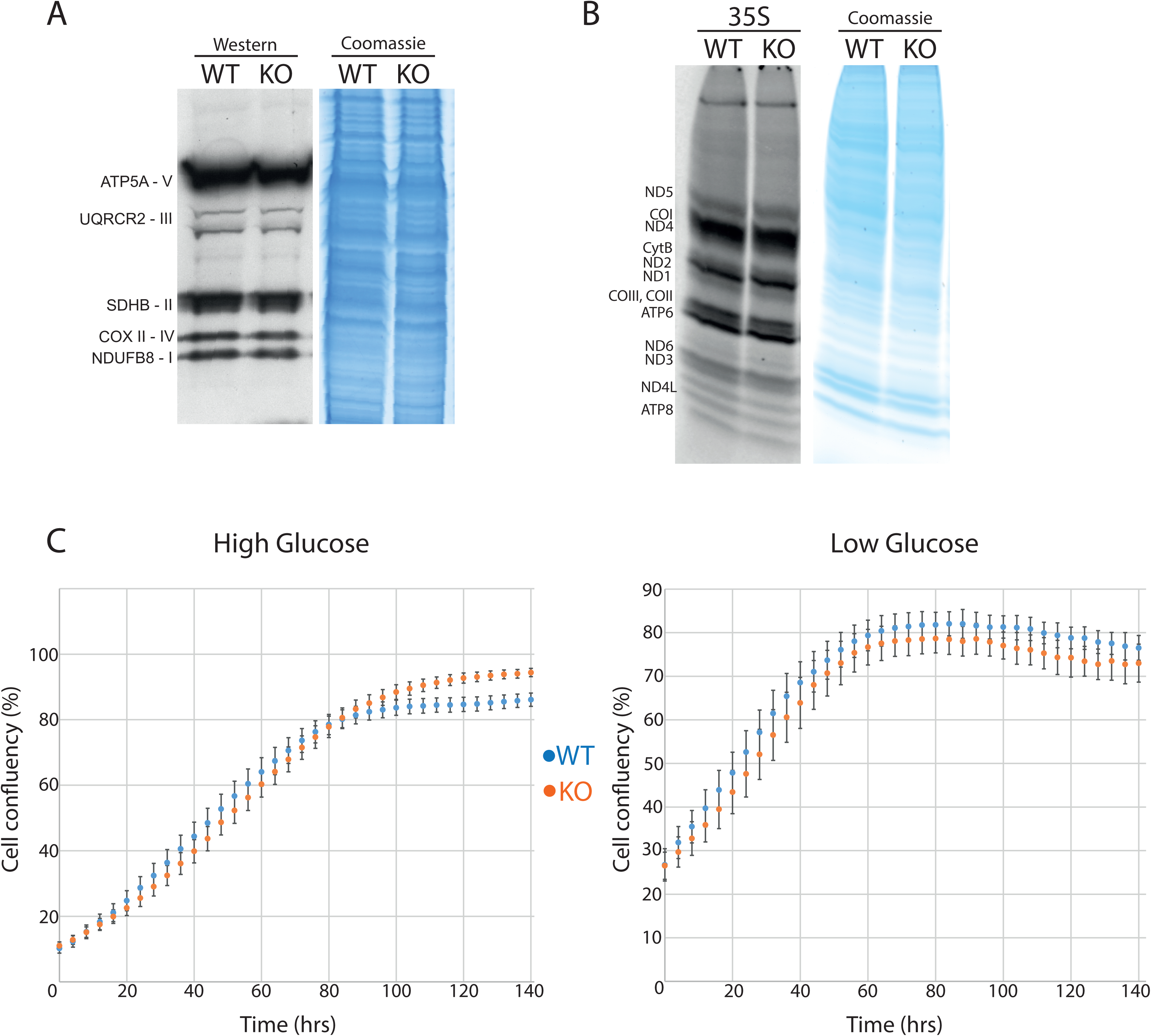
Mitochondrial translation following the loss of TRMT2B. **(a)** Western blot analysis of steady-state levels of OXPHOS subunits in HAP1 parental cell line (WT), and the HAP1 TRMT2B knockout cell line (KO). **(b)** Assessment of *de novo* mitochondrial protein synthesis through [^35^S]-methionine incorporation in the HAP1 WT and KO cell lines. Coomassie gel staining was used as a control for protein loading. **(c)**Growth curves obtained by Incucyte kinetic imaging system of HAP1 WT and KO cell lines. Cells were grown for 140 hours in the presence of ‘high’ (4.5 g/L) or ‘low’ (1 g/L) glucose.

## Discussion

### TRMT2B catalyses the formation of m5U54 and m5U429 in mt-tRNAs and 12S rRNA, respectively

In *S.cerevisiae*, the methyltransferase Trm2 is dual localised, catalysing the formation of m^5^U54 in both cyto-tRNAs and mt-tRNAs^21^, however a homology search of the human genome identified two potential orthologs, TRMT2A and TRMT2B. On the basis of mitochondrial localisation and the loss of m^5^U54 in mt-tRNAs in TRMT2B-ablated cells, this work concludes that TRMT2B is responsible for the m^5^U54-methyltransferase activity in human mitochondria. By extension, this work corroborates an earlier finding that TRMT2A is responsible for this activity in the cytosol^25^. In addition, the presence of m^5^U429 in 12S mt-rRNA, predicted from its detection in hamster mitochondria, is supported by this study, and also appears to be a product of TRMT2B. This makes TRMT2B the second human mitochondrial methyltransferase found to operate on both mt-tRNAs and mt-rRNA, following the revelation that the mt-tRNAm^1^A58-methyltransferase TRMT61B also introduces the same modification at position 947 in 16S mt-rRNA^41^. The existence of m^5^U modifications in both tRNAs and rRNAs is also well known within Archaea and Bacteria, however here they are performed by separate enzymes. For example, *E.coli* expresses three m^5^U-methyltransferase paralogs, TrmA catalysing m^5^U54 in tRNAs, RlmC catalysing m^5^U747 in 23S rRNA, and RlmD catalysing m^5^U1939 in 23S rRNA^42^.

### How does TRMT2B methylation contribute to mitochondrial translation?

The near-ubiquity in bacterial and eukaryotictRNA sequences^26^,decreased tRNA stability *in vitro*^36^, and a fundamental role in tRNA structure suggested by the convergent evolution of sterically similar m^1^Ψ54^43^, would lead one to the conclusion that m^5^U54 holds a pivotal role in translation, with severe consequences resulting from its loss. In reality, the precise biological role for this modified nucleotide has proven difficult to determine since its identification more than 50 years ago^44^. The present study identifies the enzyme responsible for this modification in human mitochondria as TRMT2B, and also shows that the same enzyme additionally catalyses the formation of m^5^U429 in 12S mt-rRNA. Furthermore, functional studies of TRMT2B appear to show the same apparent lack of phenotype identified following the deletion of yeast Trm2, with no defect observed in mt-RNA stability, mitochondrial translation, or cell growth following the loss of TRMT2B. The perceived lack of phenotype in both yeast and human cells is likely the result of either the effect being too subtle and so below the detection limits of the experiments performed, or the effect is context specific, and the environmental conditions required for the effect to become evident have not yet been applied. An effect such as this is observed for the human cyto-tRNA modifiers NSUN2 and METTL1, where phenotypes are only observed when both enzymes are depleted in conjunction^37^. A degree of functional redundancy may be a common feature of tRNA ‘core modifications’, with the absence of two or more, or their loss together with primary sequence changes, required to induce defects in translation. It is also likely that phenotypes are not being observed simply because the growth conditions are not challenged in the right manner, therefore external environmental changes such as the presence of ribosome inhibitors will also be required to differentiate between the wild type and knockout cell line.

The absence of a catalytically active TRMT2B in both bovine and rat mitochondria also raises doubts surrounding the importance of m^5^U54 in mitochondrial tRNAs relative to their cytosolic counterparts. Mitochondrial tRNAs are well known for their sometimes dramatic divergences from well conserved features found in ‘canonical tRNAs’ present in bacteria or eukaryotic cytosol^45^. An altered reliance on otherwise well conserved tRNA modifications is likely to also be a product of these significant shifts in the RNA primary sequences. The high degree of sequence conservation evident in the remainder of bovine and rat TRMT2B is not indicative of a product of gene decay. Instead, it appears likely that bovine TRMT2B retains its RNA binding properties, perhaps now acting solely as a tRNA chaperone. If further work is able to uncover a phenotype resulting from the loss of TRMT2B, it will be important to determine the extent to which it can be rescued by a catalytic mutant, in order to unravel the relative contributions made by m^5^U54 and tRNA binding.

To date, 10 modified bases have been identified in the rRNA of mammalian mitochondria; with this work establishing TRMT2B as responsible for m^5^U429, the enzymes responsible are now known for all of the modified sites determined thus far. With regard to the other modified sites found alongside m^5^U429 in the 12S rRNA, the loss of the modification has been clearly demonstrated to have a detrimental effect on mitochondrial translation. In the recent identification of METTL15 as responsible for the catalysis of m^4^C839, the loss of the enzyme was shown to impair mitoribosome assembly with a concomitant reduction in mitochondrial protein synthesis (Van Haute et al., *in press*). Likewise, loss of either NSUN4 catalysing m^5^C841^46^ and TFB1M catalysing m_2_^6^A936 and m_2_^6^A937^47^, have also been shown to impact mitoribosome assembly. Similar effects have also been observed for modifications made to the 16S rRNA. Depletion of MRM2 and MRM3, responsible for the 2’-*O*-ribose methylations Um1369 and Gm1370, respectively, results in a corresponding disruption to the assembly of the large mitoribosome subunit^48^. Likewise, diminished Ψ1397 resulting from RPUSD4 depletion has been shown to reduce the steady state levels of 16S rRNA, induce aberrant assembly of the mitoribosome large subunit, and severely reduce the rate of mitochondrial translation^49^. Future work, perhaps focusing on specific cell types, developmental stages, or environmental conditions, will be required to uncover the contribution of TRMT2B to mitochondrial translation.

## Materials and Methods

### Cell Maintenance

All cell lines were maintained in humidified incubators at 5 % CO_2_ and 37 °C, unless otherwise stated. Cells were cultured in Dulbecco’s Modified Eagle Medium (DMEM), containing 4.5 g/L glucose, 110 mg/L sodium pyruvate, supplemented with 10 % foetal bovine serum (FBS), 100 U/ml penicillin and 100 μg/ml streptomycin. Wild-type and TRMT2B Knockout (Product ID: HZGHC004364c004) HAP1 cell lines were purchased from Horizon Discovery.

### siRNA-mediated protein depletion

Stealth siRNAs (HSS129214, HSS129216, and HSS188304) were obtained from Thermo Fisher Scientific and transfected into HeLa cells using Lipofectamine RNAiMAX Reagent (Thermo Fisher Scientific).

### Subcellular Localisation by Confocal Microscopy

The full-length TRMT2B cDNA construct (Source Bioscience, CatNo: IRATp970B1220D) was cloned into a pmaxGFP vector. HeLa cells were grown on coverslips and transiently transfected with the TRMT2B-GFP construct using Lipofectamine 2000 (ThermoFisher Scientific) and after 24 hours cells were visualised by confocal microscopy, with cells treated with DAPI for nuclear staining and mitochondria immuno-stained for TOM20 as described previously^50^.

### Detection of 5-methyluridine by primer extension

RNA was extracted from cells at 60 - 80 % confluency using TRIzol reagent (ThermoFisher Scientific), following the manufacturer’s instructions. The Aniline-Hydrazine cleavage of total RNA was performed in a two-step procedure: the incubation of RNA with hydrazine which modifies uridines and its derivatives, followed by the cleavage of hydrazine-modified RNA at uridines by aniline. The resulting cleavage is then detected via primer extension (**Figure S2**). To begin, 5 μg of pelleted RNA was resuspended in 20 μL of chilled 50 % hydrazine hydrate and incubated on ice for 5 minutes. The reaction was then stopped with the addition of a 1/10th volume of 3 M NaOAc at pH 5.2 and 3 volumes of 100 % ethanol. The sample was then precipitated at -80 °C for>12 hours. Following this, the precipitated hydrazine-treated sample was centrifuged and the pellet resuspended in 1 M aniline-acetate pH 4.5. The sample was incubated at 60 °C for 20 minutes, and the RNA then precipitated again and resuspended in 30 μL of water.

The reverse transcription primer extension reactions were performed as in (^48^), with the following primers used:

mt_Pro_m^5^U54: TGGTCAGAGAAAAAGTC

mt_12S_m5U429: GGGCTAAGCATAGTGGGGTATC

cyto-Asp_m5U54: TGGCTCCCCGTCGGGGAATC

cyto-Glu_m5U54: TGGTTCCCTGACCGGGAAT

### Northern Blotting

RNA was extracted from cells at 60 - 80 % confluency using TRIzol reagent (ThermoFisher Scientific) and subjected to northern blotting as described previously^51,52^. Briefly, total RNA was resolved on 1% agarose gels containing 0.7?M formaldehyde in 1× MOPS buffer (mt-rRNAs) or on 10% UREA–PAGE in 1× GTB buffer (mt-tRNAs), or on 6.5% UREA–PAGE in 100mM NaOAc pH 5.0 (aminoacyl-mt-tRNAs), transferred to a nylon membrane, ultraviolet-crosslinked to the membrane and hybridized with radioactively labelled T7-transcribed radioactive RNA-probes.

### Western Blotting

20-30 µg of total extracted protein was diluted to an equal volume and a ⅓ volume of NuPAGE LDS 4 x sample buffer and loaded on SDS-PAGE 4-12% bis-tris gels (Life Technologies) and transferred onto a membrane using iBlot 2 Dry Blotting System (Thermo Fisher Scientific). The following antibodies were used: rabbit anti-MRPS17 (Proteintech 18881-1-AP, 1:1000), rabbit anti-MRPS18b (Proteintech 16139-1-AP, 1:1000), rabbit anti-MRPS35 (Proteintech 16457-1-AP, 1:1000), Total OXPHOS Human WB antibody cocktail (Abcam, ab110411, 1:1000), goat anti-rabbit IgG HRP (Promega W4011, 1:2000), goat, anti-mouse IgG HRP (Promega W4021).

### [^35^ S]-methionine metabolic labeling of mitochondrial proteins

The labelling of newly synthesised mtDNA-encoded proteins was performed as in Rorbach *et al*.^48^. Briefly, exponentially growing cells were incubated in methionine/cysteine-free medium for 10 min before the medium was replaced with methionine/cysteine-free medium containing 10% dialysed FCS and emetine dihydrochloride (100 µg/mL) to inhibit cytosolic translation. Following a 20 min incubation, 120 μCi/mL of [^35^S]-methionine (Perkin Elmer) was added and the cells were incubated for 30 min. Cells were washed with PBS, lysed, and 30µg of protein was loaded on 10-20% Tris-Glycine SDS-PAGE gels. Dried gels were visualized and quantified with a PhosphorImager system with ImageQuant software.

## Supplementary material

Supplementary data for this article can be accessed [link]

## Acknowledgements

We would like to thank the members of the Mitochondrial Genetics Group at the MRC-MBU for stimulating discussions during the course of this work. Maciej Szukszto is acknowledged for his technical assistance in the experiments presented in Figure 2.

## Authors’ contribution

C.A.P. conceived, designed and performed the experiments, undertook the data analysis and wrote the manuscript; M.M. oversaw the project and wrote the manuscript. Both authors commented and approved the final version of the manuscript.

## SUPPLEMENTAL FIGURES

**Figure S1:**
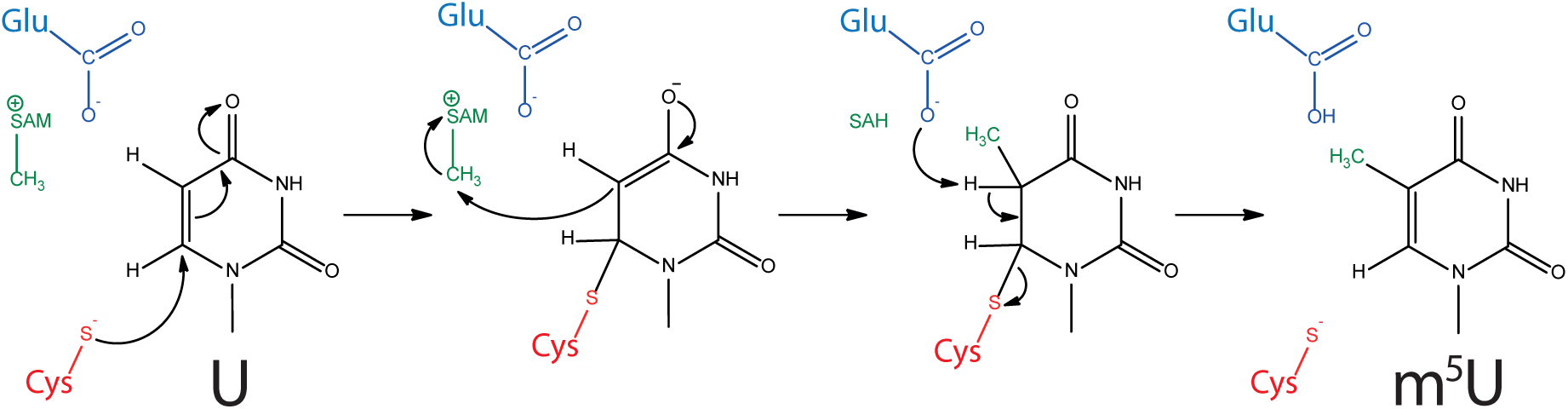
The catalytic mechanism of m^5^U-methyltransferases, showing the involvement of the nucleophilic cysteine, the proton extracting glutamate, and S-adenosylmethionine (SAM).

**Figure S2:**
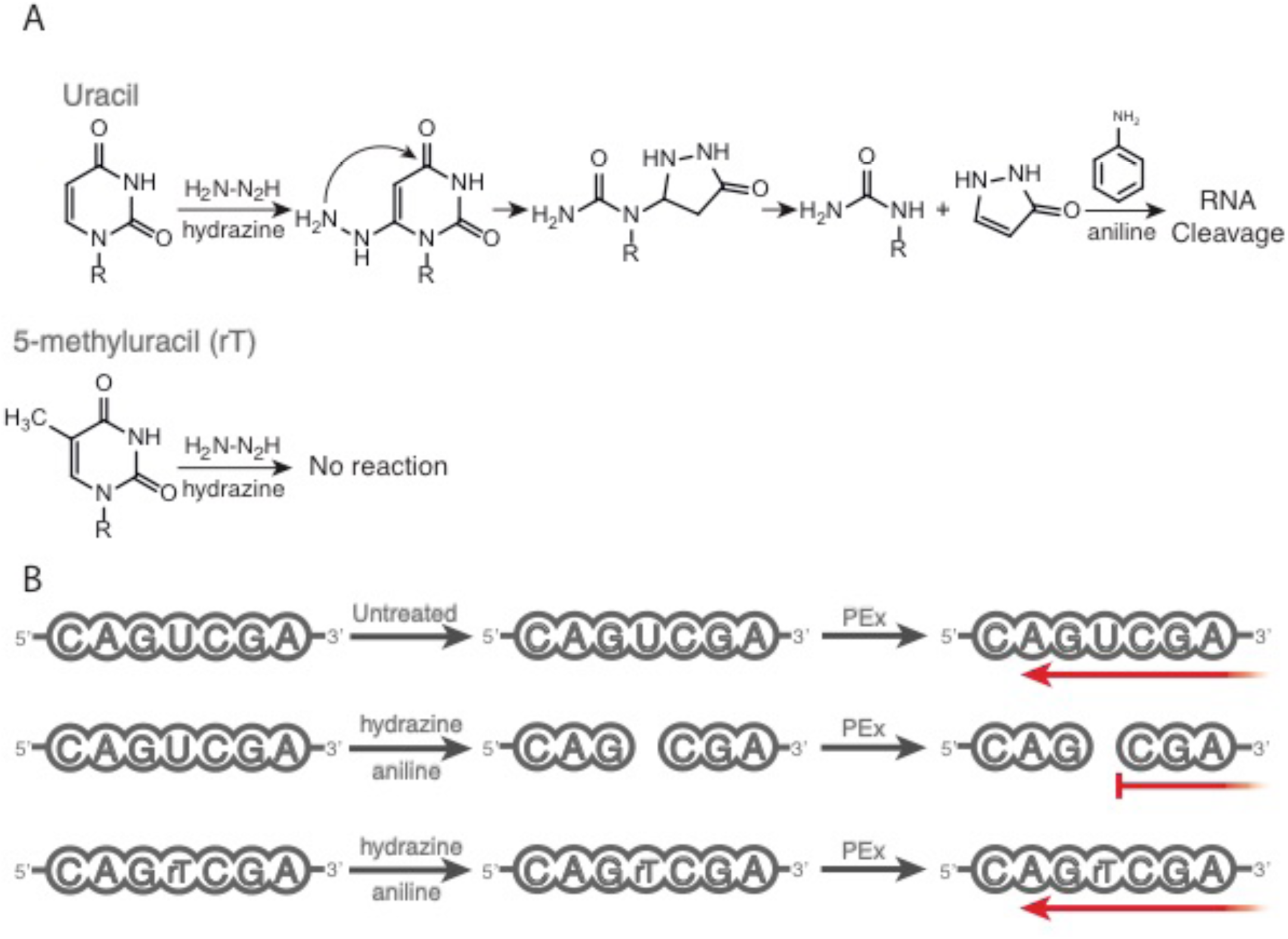
Detection of 5-methyluridine in RNA. **(a)** The hydrazinolysis of uracil in RNA is mediated through the nucleophilic attack of C6 by hydrazine, and the subsequent reaction with C4, forming a pyrazole which is spontaneously cleaved from the glycosidic nitrogen. The resulting abasic site is susceptible to β-elimination at both the 3’ and 5’ phosphates in the presence of aniline. A methyl group at C5 protects the pyrimidine ring from nucleophilic attack, and therefore subsequent strand cleavage at this position **(b)** Primer extension (red arrow) following Hydrazine-Aniline treatment will result in stalled extension products corresponding to unmodified uracils in the original sequence, whereas primers will read through m^5^U. These products are then separated through gel electrophoresis.

**Figure S3:**
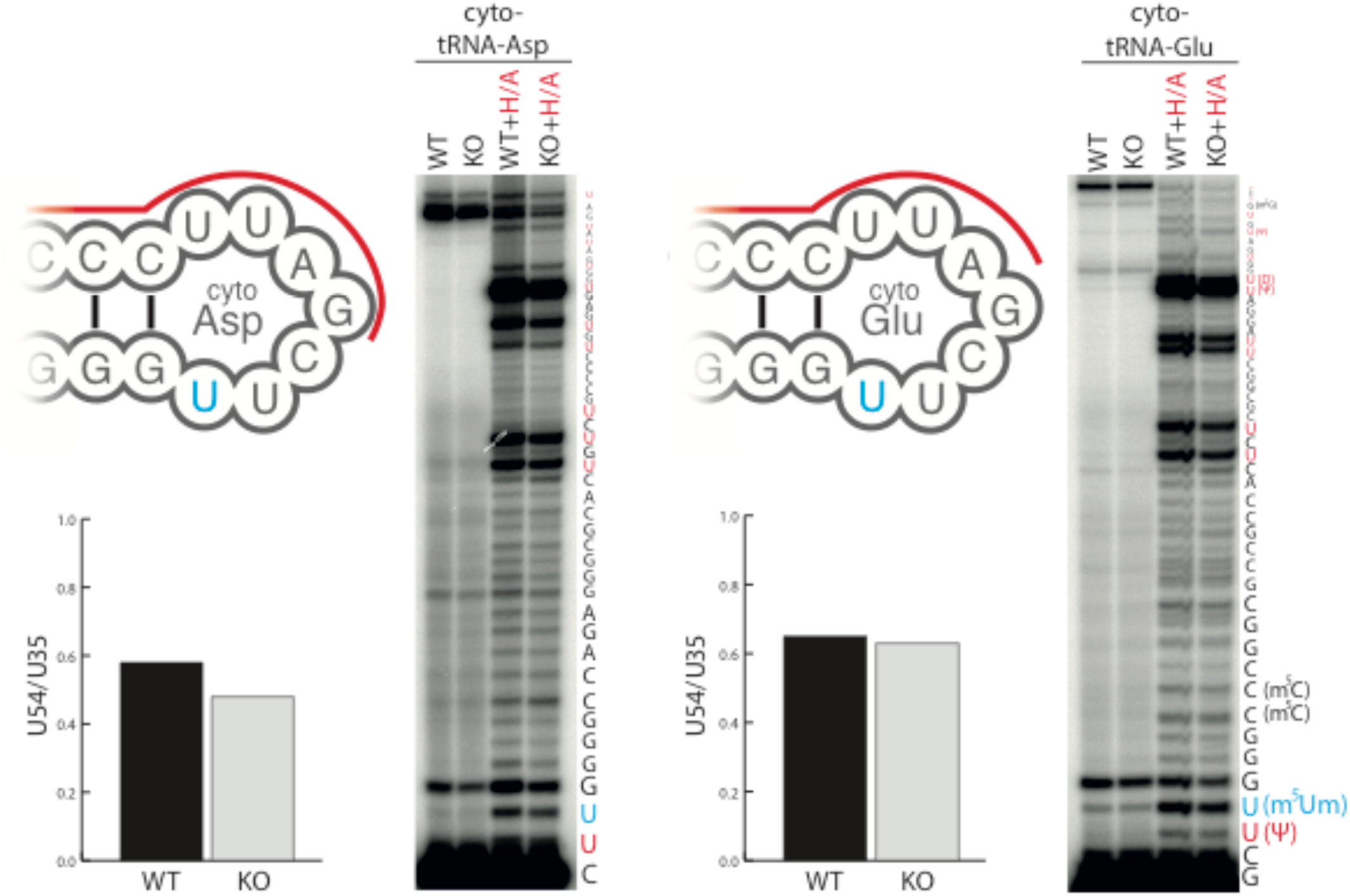
m^5^U54 in cyto-tRNAs is retained in TRMT2B KO cells. Separation and detection of RT-PEx products using a [^32^P]-end labelled primer complementary to the region upstream of m^5^U54 (red line) in cyto-tRNAAsp and cyto-tRNAGlu. Extension reactions performed on RNA derived from a HAP1 wild-type (WT) TRMT2B knockout cell line (KO), and following hydrazine-aniline treatment (WT+H/A and KO + H/A). Quantification values represent the ratio between stalling at U54 and stalling at the next uridine residue (U35), after the values in the untreated lanes had been subtracted from both to account for background. The nucleotide sequence of the tRNA, corresponding to stalling events at each position, is shown to the side of the panel, with the addition of the modifications present for cyto-tRNA^Glu^ where the modification profile is known.

